# Chrysalis: A new method for high-throughput histo-cytometry analysis of images and movies

**DOI:** 10.1101/403154

**Authors:** Dmitri I. Kotov, Thomas Pengo, Jason S. Mitchell, Matthew J. Gastinger, Marc K. Jenkins

## Abstract

Advances in imaging have led to the development of powerful multispectral, quantitative imaging techniques, like histo-cytometry. The utility of this approach is limited, however, by the need for time consuming manual image analysis. We therefore developed the software Chrysalis and a group of Imaris Xtensions to automate this process. The resulting automation allowed for high-throughput histo-cytometry analysis of 3D confocal microscopy and two-photon time-lapse images of T cell-dendritic cell interactions in the spleen. It was also applied to epi-fluorescence images to quantify T cell localization within splenic tissue by using a ‘signal absorption’ strategy that avoids computationally intensive distance measurements. In summary, this image processing and analysis software makes histo-cytometry more useful for immunology applications by automating image analysis.

## Introduction

Imaging of biological samples has traditionally been used to resolve anatomic structures (1) or identify specific cells in tissues (2). Recent advances in image analysis, like histo-cytometry (3) and dynamic in situ cytometry (4) have expanded the depth of analysis by increasing characterization of cell types and objective quantification of cells in images. These new techniques combine multispectral image analysis with a quantitative workflow. The image quantification is performed by analyzing image-derived statistics in flow cytometry analysis software (3, 4). These approaches can quantify the number and location of cells throughout a tissue (5), identify cell-cell interactions (6), and correlate protein expression to cellular localization (7). Histo-cytometry and dynamic in situ cytometry have been applied to a variety of imaging systems including confocal (8–10), epi-fluorescence (11, 12), and two-photon microscopy (4). However, these approaches are time consuming due to the need for extensive hands-on image processing. We addressed this issue by creating the software Chrysalis and a suite of Imaris Xtensions to batch image processing and analysis (https://histo-cytometry.github.io/Chrysalis/). This automation reduced hands-on analysis time for confocal, epi-fluorescence, and two-photon microscopy images. The broad applicability of this protocol was confirmed by quantifying cell localization and cell-cell interactions in the spleen using multiple imaging platforms. Automation should facilitate the use of the powerful histo-cytometry technique.

## Materials and Methods

### Mice

Six-to eight-wk old C57BL/6 (B6) female mice were purchased from the Jackson Laboratory or the National Cancer Institute Mouse Repository (Frederick, MD, USA). *Itgax*^*YFP*^ (13) and *Rag1*^*-/-*^ *Ubc*^*GFP*^ (14) TEa TCR transgenic (15) female mice were a gift from B.T. Fife (University of Minnesota). *Rag1*^*-/-*^ B3K506 TCR transgenic (16) and *Rag1*^*-/-*^ B3K508 TCR transgenic mice (16) were bred and housed in specific pathogen–free conditions in accordance with guidelines of the University Institutional Animal Care and Use Committee and National Institutes of Health. The University Institutional Animal Care and Use Committee approved all animal experiments.

### Infections

Mice were injected i.v. with 10^7^ colony-forming units of ActA-deficient *Listeria monocytogenes* (*Lm*) expressing the P5R peptide (*Lm*-P5R) (17, 18).

### Cell transfer

Lymph nodes were collected from *Rag1*^*-/-*^ B3K506 TCR transgenic, *Rag1*^*-/-*^ B3K508 TCR transgenic, and *Rag1*^*-/-*^ *Ubc*^*GFP*^ TEa TCR transgenic mice and a small sample was stained with APC-labeled CD4 antibody (RM4-5, Tonbo biosciences) and analyzed on an LSR II (BD Biosciences) flow cytometer using Flowjo software (TreeStar). The results were used to calculate the amount of the remaining sample needed to transfer one million CD4^+^ T cells. In some cases, the T cells from the *Rag1*^*-/-*^ B3K506 and *Rag1*^*-/-*^ B3K508 TCR transgenic mice were also labelled with CellTracker Orange (ThermoFisher Scientific) or CellTraceViolet (ThermoFisher Scientific), respectively (19). One million TCR transgenic cells were transferred into B6 mice by i.v injection 24 h prior to infection with *Lm*-P5R.

### Confocal microscopy

Twenty µm splenic sections from naive or *Lm*-P5R infected mice were stained with Brilliant Violet (BV) 421-conjugated F4/80 (BM8, Biolegend), Pacific Blue-conjugated B220 (RA3-6B2, Biolegend), CF405L-conjugated CD8α (53-6.7, Biolegend), AF488-conjugated pS6 (2F9,Cell Signaling Technologies), CF555-conjugated CD86 (GL-1, Biolegend), AF647-conjugated CD45.2 (104, Biolegend), AF700-conjugated MHC II (M5/114.15.2, Biolegend), CF514-conjugated CD11c (N418, Biolegend), BV480-conjugated CD3 (17A2, BD biosciences), and AF594-conjugated SIRPα (P84, Biolegend) antibodies. Certain purified antibodies from Biolegend were conjugated with CF405L, CF514, or CF555 with Biotium Mix-n-Stain labelling kits. Confocal microscopy was performed with a Leica SP5 confocal microscope with two HyD detectors; two PMT detectors; 405, 458, 488, 514, 543, 594 and 633 laser lines; and a 63X oil objective with a 1.4 numerical aperture. The mark and find feature in the Leica Application Suite was used to image 12 T cell zones in each spleen with each image consisting of a 20 µm z-stack acquired at a 0.5 µm step size. Additionally, the Leica SP5 microscope was used to image single color stained Ultracomp eBeads (ThermoFisher Scientific) for generating a compensation matrix.

### Epi-fluorescence microscopy

Spleens from B6 mice infected 48 h earlier with *Lm*-P5R were fixed with paraformaldehyde, dehydrated with sucrose, and embedded in OCT. Seven µm sections of these spleens were stained with BV421-conjugated F4/80, AF488-conjugated B220 (RA3-6B2, Biolegend), AF647-conjugated CD45.2 (104, Biolegend), and AF594-conjugated CD3 (17A2, Biolegend) antibodies. The samples were imaged with a Leica DM6000B epi-fluorescence microscope equipped with a dry 20X objective with 0.5 numerical aperture and a Leica DFC 9000 camera with custom filter cubes. The tiling feature in the Leica Application Suite (Leica Microsystems) software was used to image the entire splenic section and the images were analyzed by following the histo-cytometry protocol (Supplemental Protoc. 1).

### Two-Photon microscopy

*Rag1*^*-/-*^ *Ubc*^*GFP*^ TEa TCR transgenic CD4^+^ T cells, CMTMR-labelled B3K506 TCR transgenic T cells, and CTV labelled B3K508 TCR transgenic T cells were transferred into *Itgax*^*YFP*^ mice that were then infected with *Lm*-P5R bacteria 24 h after cell transfer. Recipient spleens were immobilized on plastic coverslips, sliced longitudinally with a vibratome, and perfused with 37° C DMEM media bubbled with 95% O_2_ and 5% CO_2_. Samples were imaged with a 4-channel Leica TCS MP microscope with a resonant-scanner containing two NDD-and two HyD-photomultiplier tubes operating at video rate. The objective was a water dipping 25X with 0.95 numerical aperture. Samples were excited with a MaiTai TiSaphire DeepSee HP laser (15 W; Spectra-Physics) at 870 nm, and emissions at 440-480 (CTV), 500–520 (GFP), 520-560 (YFP) and 560-630 (CMTMR) nm were collected. Images acquired were 20–250 µm below the cut surface of the spleen slice and 512×512 XY frames were collected at 3.0 µm steps every 30 s for 30 min.

### Image processing and histo-cytometry analysis

A compensation matrix was created in ImageJ (NIH) by using the GenerateCompensationMatrix script on images of single color stained Ultracomp eBeads. This compensation matrix was applied to 3D images and movies in Chrysalis to compensate for the spillover of each fluorescent signal from its channel into other channels. Chrysalis was also used for further automated image processing as described in Fig. 1A and Fig. 5A. Imaris 8.3, 8.4, 9.0, and 9.1 (Bitplane) were used for image analysis, including surface creation to identify cells in images. The Sortomato V2.0, XTChrysalis, and XTChrysalis2phtn Xtensions were used in Imaris for identifying cellular subsets based on protein expression, quantifying cell-cell interactions, and exporting cell surface statistics. Statistics were exported from these applications and imported into FlowJo v10.3 (Treestar) for quantitative image analysis. Details of these steps are included in Supplemental Protoc. 1.

**Figure 1.**
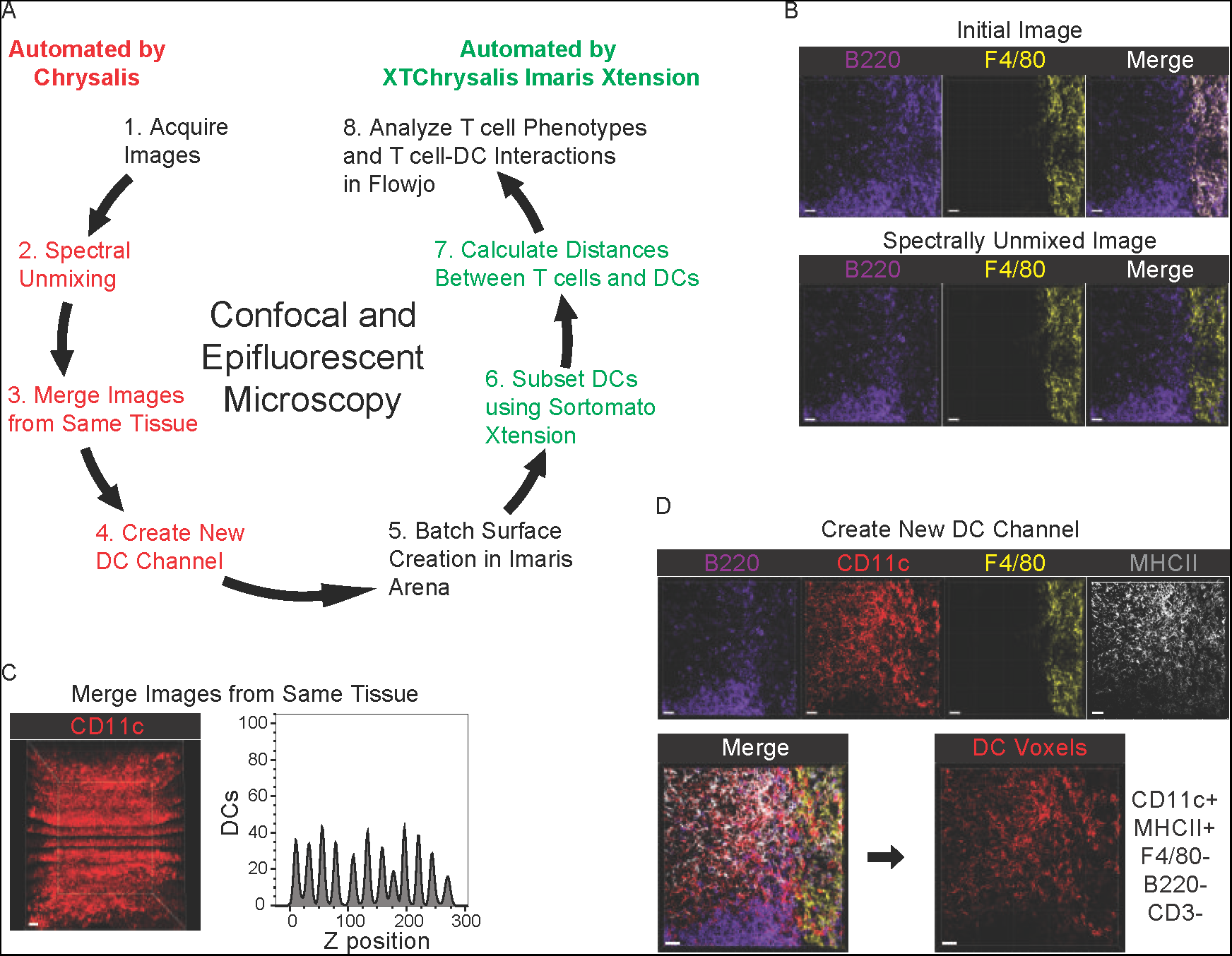
Image Processing with Chrysalis. (A) Diagram of the histo-cytometry workflow on 3D images when automated by Chrysalis and XTChrysalis. (B) B220 and F4/80 staining of splenic tissue before and after spectral unmixing in Chrysalis. (C) CD11c staining and histogram of DCs in 12 confocal microscopy images merged together in the z plane. (D) Generation of a DC voxel channel with Chrysalis’ new channel feature by utilizing the fluorescence of existing channels including B220, CD11c, F4/80 and MHCII, which are depicted for a splenic tissue section. Scale bar, 20 µm. Data representing two to three independent experiments are shown.

### Code availability

All of the code generated for image processing or analysis can be downloaded at https://histo-cytometry.github.io/Chrysalis/, including compiled versions of Chrysalis for Windows and Mac OSX with a Linux version available upon request due to Github limitations on file size. Additionally, all of the Imaris Xtensions are compatible with Windows and Mac OSX. The documentation for the code as well as a detailed protocol for image acquisition and analysis is provided in Supplemental Protoc. 1.

## Results

### Automated Processing of 3D images

Image acquisition, processing, and analysis with histo-cytometry consists of eight steps (Fig. 1A). We developed a stand-alone software called Chrysalis for automating the three image processing steps (steps 2-4) as well as a suite of Imaris Xtensions that automate two of the image analysis steps (steps 6 and 7; Fig. 1A). For processing 3D images, Chrysalis spectrally unmixes images, merges images, and generates new channels prior to image analysis in Imaris (Fig. 1A). Each of these features addresses existing issues with standard image analysis workflows and expedites image analysis. For example, spectral unmixing accounts for spectral overlap between different fluorophores and fluorescent proteins (20). To aid in this step, we wrote a script that automatically generates a compensation matrix from user-provided single-color control images. Chrysalis uses this compensation matrix to spectrally unmix an image with a linear unmixing algorithm (Fig. 1B) (21).

Another issue addressed by Chrysalis is the image processing required for efficiently analyzing cell-cell interactions in 3D images. When analyzing cell-cell interactions, high magnification images need to be taken to observe the interaction event. Analysis of interactions in large tissues like spleen or lymph node can be performed by tiling images of the entire tissue together. However, this process is extremely time intensive for image acquisition and analysis due to the high magnification and large number of images required. This approach is also inefficient in cases where the interaction event occurs only in a small percentage of the tissue. Rare interactions within 3D images can instead be identified at the microscope allowing for the acquisition of only the images that depict the relevant interactions at high magnification prior to manually merging the images together for analysis. Such a process was previously applied to analyze T regulatory cell-dendritic cell (DC) clusters (7). To make it easier to study rare interaction events, Chrysalis can automatically merge multiple images from one tissue through stacking images in the z plane (Fig. 1C), which allows for time-efficient and consistent analysis of the relevant cell-cell interaction event.

Some cell types require identification based on expression of multiple proteins. For example, DCs are identified by their expression of CD11c and MHC class II (MHCII), but not B220, F4/80, or CD3 (13, 22, 23). To address this issue, Chrysalis creates new channels consisting of voxels that are above a computer-generated threshold (24) for user-selected “include” channels and below a computer-generated threshold for user-selected “exclude” channels, a process called voxel gating (3). A user-selected base channel expressed by the cell type dictates the signal intensity in this new channel. For a new DC channel, CD11c and MHCII would be the include channels, while B220, F4/80, and CD3 would be the exclude channel and the base channel would be CD11c (Fig. 1D). In effect, this new channel provides better DC resolution than the CD11c channel alone.

### Automated histo-cytometry analysis of 3D images

For histo-cytometry analysis, Chrysalis processed images are imported into the image analysis software Imaris, which creates surfaces to identify cells based on the image’s channels (3, 8, 10, 25). Traditionally, the steps required to analyze surfaces requires extensive hands-on time. Thus, we created the Xtension XTChrysalis, which automates this process. XTChrysalis 1) separates existing surfaces into new surfaces based on a gating scheme defined in a Xtension called Sortomato, 2) calculates distances to each new surface, 3) rescales signal intensities for any images, and 4) exports statistics for any surface (Fig. 1A). The exported statistics contains each channel’s intensity mean and minimum values as well as each cell’s volume, sphericity, and position. All values have 0.1 added to them to enable logarithmic display of each parameter. This data can be directly imported into quantitative analysis software, like Flowjo or XiT (26), for further analysis.

### Analyzing T cell activation and T cell-DC interactions in 3D images

To demonstrate 3D image analysis with Chrysalis and XTChrysalis, we analyzed images of T cells, DCs, and their interactions captured by confocal microscopy. Following infection, DCs interact with T cells by presenting MHCII-bound peptides derived from the invading pathogen, leading to TCR signaling (27, 28). To analyze this type of interaction, splenic tissue from *Lm* infected mice was analyzed by 10 color confocal microscopy. T cell responses were examined using a system involving adoptive transfer of B3K506 TCR transgenic (Tg) CD4^+^ T cells that express P5R peptide:MHCII-specific TCRs. B3K506 TCR Tg T cells were injected into B6 recipient that were then infected with *Lm*-P5R bacteria. Twenty-four h after infection, 12 T cell zones were imaged per spleen to obtain sufficient cells for analysis (29). We used Chrysalis to spectrally unmix, rescale, and merge images, and generate a new channel representing DC voxels before image analysis in Imaris (Fig. 2A). TCR Tg cell surfaces were then created based on CD45.2 fluorescence (Fig. 2B). Staining for the phosphorylated form of S6 kinase (pS6), an indicator of TCR signaling (30), was examined within those surfaces to identify cells undergoing TCR signaling (Fig. 2B). DC surfaces were generated based on the DC voxel channel, thereby identifying hundreds of DCs (Fig. 2C). The Sortomato Xtension was used to identify a gating strategy to subset the DCs based on expression of CD8α or SIRPα (Fig. 2D) (22, 31–33). XTChrysalis was then applied to the processed images and the resulting data was analyzed in Flowjo.

**Figure 2.**
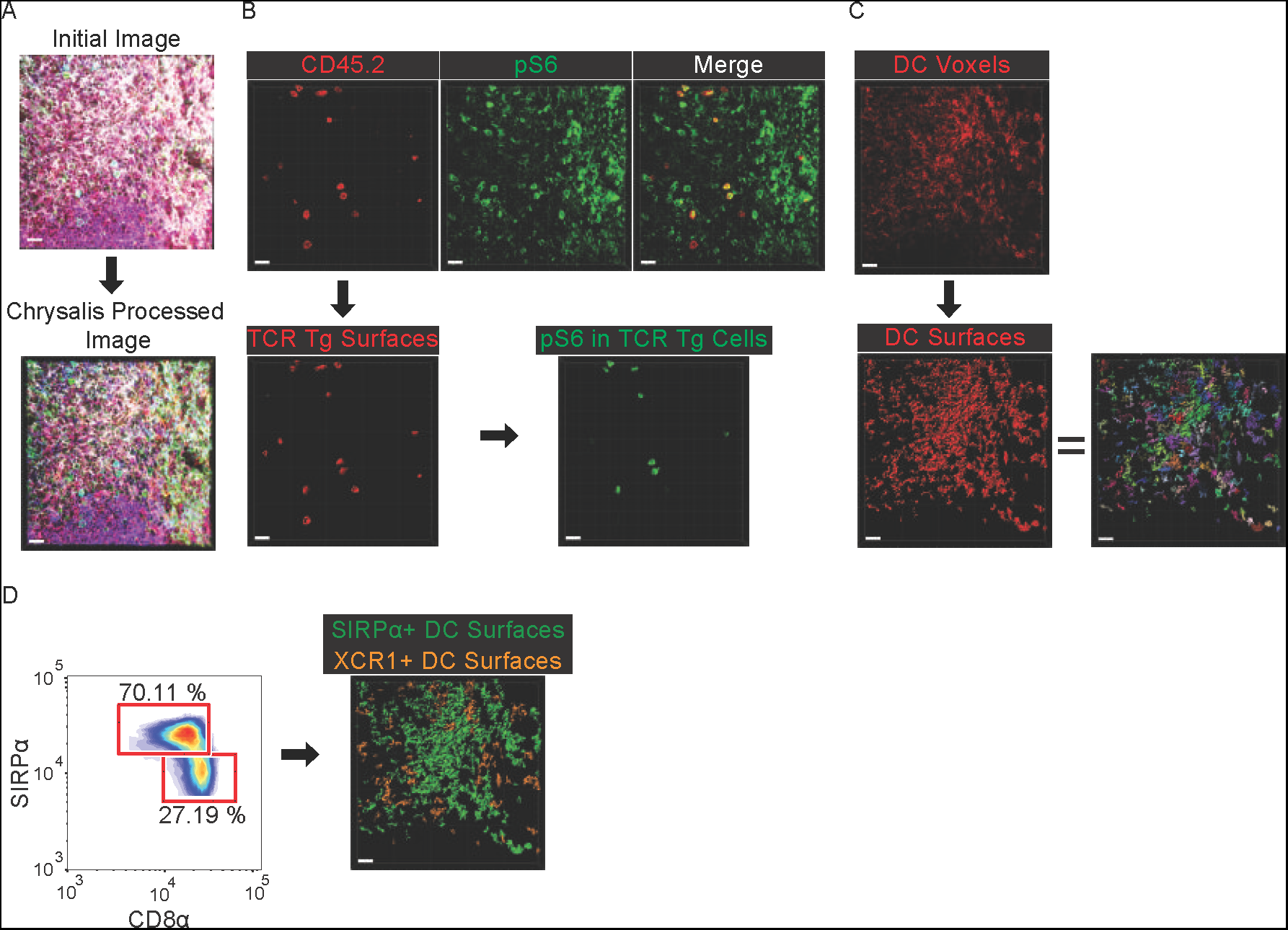
Chrysalis and XTChrysalis analysis of a 3D image. (A) Confocal microscopy 10 color image before and after Chrysalis processing. (B) Identifying TCR Tg cells with CD45.2 staining and TCR signaling based on pS6 expression. (C) DC voxels (CD11c^+^ MHCII^+^ B220^-^ CD3^-^ F4/80^-^) that were used to identify DCs by surface creation in Imaris. (D) 2D plot generated with Sortomato for subsetting DC surfaces into SIRPα^+^ or XCR1^+^ DCs based on SIRPα and CD8α expression. Scale bar, 20 µm. Data representing two to three independent experiments are shown.

As expected, B3K506 TCR Tg cells expressed CD45.2, and DCs expressed CD11c and MHCII (Fig. 3A). Surprisingly, there were two populations of TCR Tg cells, one lacking CD11c and MHCII signals and one with these signals (Fig. 3B). The populations were similar in cell size but differed in TCR signaling with the MHCII^high^ CD11c^high^ population having greater TCR signaling based on pS6 expression (Fig. 3B). Since MHCII and CD11c are not expressed by T cells (34), we hypothesized that the TCR Tg cell surfaces “absorbed” the DCs’ MHCII and CD11c signals by being in close proximity to DCs. This hypothesis was tested by comparing the frequency of T cell-DC interactions for the MHCII^high^ CD11c^high^ and the MHCII^low^ CD11c^low^ T cells. The MHCII^high^ CD11c^high^ T cells interacted with XCR1^+^ and SIRPα^+^ DCs 10 times as often as the MHCII^low^ CD11c^low^ T cells, suggesting that the DC signal absorption hypothesis was correct (Fig. 3C).

**Figure 3.**
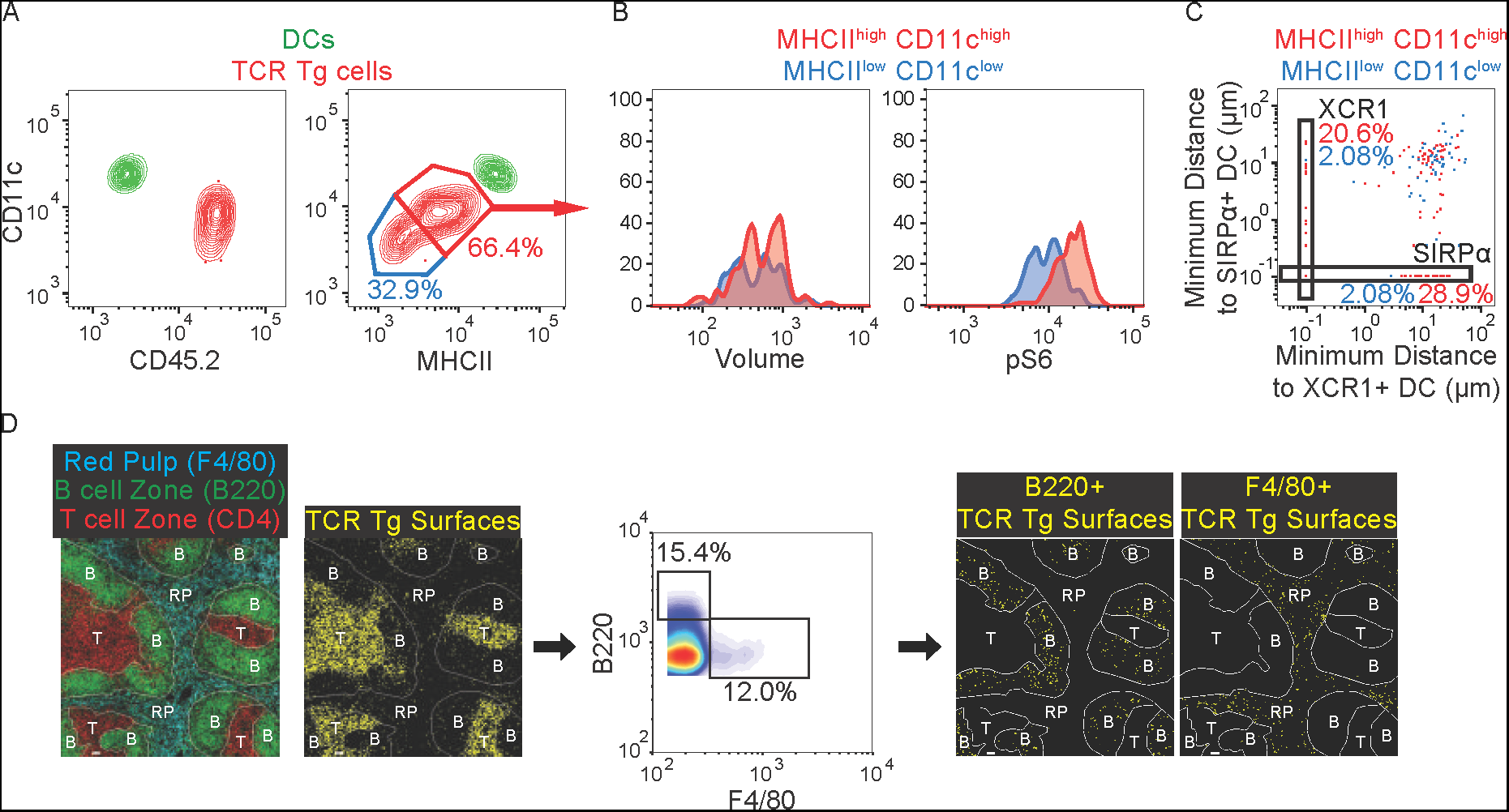
The ‘signal absorption’ strategy can accurately quantify cell-cell interactions and cellular localization. (A) Flowjo analysis of CD11c, CD45.2, and MHCII expression on DCs (green) and B3K506 TCR Tg T cells (red) identified in confocal microscopy images. (B) Histogram of volume and pS6 expression for MHCII^high^ CD11c^high^ (red) and MHCII^low^ CD11c^low^ (blue) TCR Tg T cells. (C) Quantifying T cell-DC interactions for MHCII^high^ CD11c^high^ (red) and MHCII^low^ CD11c^low^ (blue) TCR Tg T cells with SIRPα^+^ and XCR1^+^ DCs. (D) Epi-fluorescence image of splenic tissue stained for F4/80, B220, CD4, and CD45.2, with TCR Tg cell surfaces created based on CD45.2 fluorescence. TCR Tg surfaces were subset into cells that absorbed B220 or F4/80 surfaces thereby allowing for the characterization of TCR Tg cell localization. Scale bar, 20 µm. Data representing two to three independent experiments are shown.

### Quantifying cellular localization in epi-fluorescence microscopy images

The experimental approach described above was also used to assess the locations of B3K506 TCR Tg cells by epi-fluorescence microscopy. Spleens from B6 recipients of B3K506 T cells infected three d earlier with *Lm*-P5R bacteria were stained for F4/80, B220, and CD4 to identify the red pulp, B cell zones, and T cell zones, respectively (Fig. 3D) (35, 36). Spleens were also stained for CD45.2 to identify the TCR Tg cells. Macrophages in the red pulp express F4/80 (36) and B cells in the B cell zone express B220 (37), but neither protein is expressed by T cells (38–40). Therefore, TCR Tg surfaces that have B220 signal should be in close proximity to B cells and reside in B cell follicles while those with F4/80 signal should be near macrophages and localize to the red pulp. Indeed, although most of the B3K506 T cells were in the T cell zones, some were in the B cell follicles and absorbed B220 signal, while others were in the red pulp and absorbed F4/80 signal (Fig. 3D). Thus, the location of a cell can be determined based ‘absorption’ of fluorescent signal from proteins expressed by nearby cells.

### The effect of TCR affinity on T cell localization

The capacity of the ‘signal absorption’ strategy to identify cell location was also employed to validate the concept that TCR signal strength influences Th cell differentiation (41). It has been shown that naïve T cells with high TCR affinity for peptide:MHCII tend to differentiate into Type 1 helper (Th1) cells while cells with lower affinity TCRs primarily adopt the T follicular helper (Tfh) fate (17, 42). These differences in T cell differentiation would be expected to modulate T cell localization because different Th subsets express different chemokine receptors. For example, Th1 cells express CXCR3 (43, 44) driving them towards sites of inflammation such as the splenic red pulp, while Tfh cells express CXCR5 allowing them to traffic into B cell follicles (45, 46). Thus, Tfh-biased low TCR affinity T cells would localize to B cell follicles at a higher frequency than Th1-biased high TCR affinity T cells.

B3K506 T cells were compared to B3K508 TCR Tg T cells, which express TCRs with lower affinity for P5R:I-A^b^ complexes, to test this hypothesis (16, 47). The TCR Tg populations were transferred into B6 mice, which were infected with *Lm*-P5R bacteria. Spleen sections were stained, imaged by epi-fluorescence microscopy, and analyzed with Chrysalis one, two, and three d after infection. As in the previous experiment (Fig. 3D), B220 identified B cell follicles, CD4 defined T cell zones, F4/80 delineated red pulp, and CD45.2 specified TCR Tg T cells (Fig. 4A). The signal absorption strategy was further validated by comparing the T cell distance into the B cell follicles with their absorption of B220. All of the T cells that were located in the B cell follicle where B220^high^ further validating the signal absorption strategy (Fig. 4B). T cell localization in the follicles or red pulp was, therefore, identified based on T cell absorption of B220 or F4/80 signal, respectively (Fig. 4C). As expected, TCR Tg cells were primarily situated in T cell zones in naive mice and during the initial three d following *Lm*-P5R infection (Fig. 4D). However, the signal absorption assay revealed a greater proportion of low TCR affinity B3K508 T cells localized to B cell follicles than high TCR affinity B3K506 T cells, in line with B3K508 T cells favoring the B cell follicle-homing Tfh cell fate (Fig. 4D) (17). This result demonstrates the ability of the improved histo-cytometry workflow to quantify cellular localization in epi-fluorescence microscopy images with a novel signal absorption strategy.

**Figure 4.**
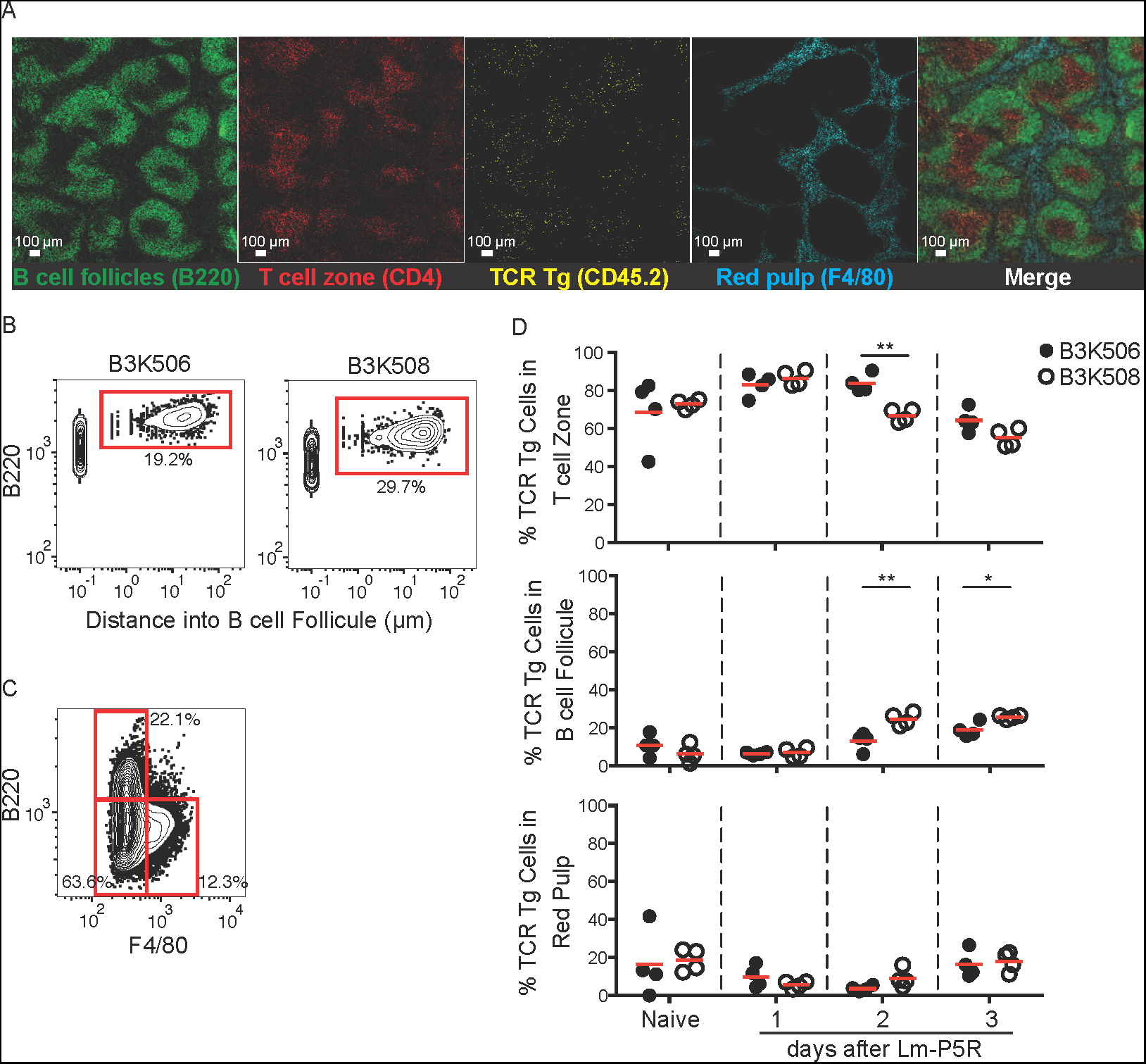
T cells primarily reside in T cell zones following *Listeria* infection, and low affinity T cells traffic into B cell follicles more than high affinity T cells. (A) Representative images of B220, CD4, CD45.2, and F4/80 staining of a splenic tissue section acquired by epi-fluorescence microscopy. (B) Flowjo analysis of the relationship between B220 absorption and T cell distance into B cell follicles for B3K506 and B3K508 cells three d after *Lm*-P5R infection. (C) Gating strategy for using ‘signal absorption’ to identify B cell follicle (B220^+^) or red pulp (F4/80^+^) residing TCR Tg T cells in epi-fluorescence microscopy images. (D) Quantification of epi-fluorescence microscopy images that determine B3K506 (filled circle, n=4) and B3K508 (empty circle, n=4) cell localization in T cell zone, B cell follicle, or red pulp in spleens of naïve mice and mice one, two, or, three d after *Lm*-P5R infection. Scale bar, 100 µm. Pooled data from three independent experiments are shown. One-way ANOVA was used to determine significance for D. * = p < 0.05, ** = p < 0.01.

### Automated processing and histo-cytometry analysis of two-photon microscopy images

Previously, histo-cytometry has been applied to 3D images, however this same methodology can be applied to two-photon time-lapse data (movies) (7, 8). Chrysalis can aid in this application because it can spectrally unmix, generate new channels, and rescale movies (Fig. 5A). Additionally, Chrysalis expedites two-photon movie analysis by simplifying existing workflows. For example, two-photon movies can have variable image quality due to poor tissue health stemming from a lack of oxygenation or low tissue temperature (48, 49). Tissue health can be assessed by examining the motility of a control population within the tissue, like fluorescently labeled polyclonal T cells (50). By reviewing the motility of a control cell population across several movies, movies that depict healthy tissue can be identified prior to conducting in-depth analysis. To optimize this process, Chrysalis processes movies by Gaussian filtering and rescales each channel to maximize signal intensity and movie clarity. The processed movies are saved as AVI files, which can be quickly examined for tissue health prior to performing more time consuming analysis.

We have also written an Imaris Xtension called XTChrysalis2phtn that batches histo-cytometry analysis of two-photon movies. For each movie, XTChrysalis2phtn will 1) calculate distances between cell surfaces and define cell-cell interactions at each time point, 2) rescale signal intensities, and 3) export statistics for each surface (e.g. average velocity, displacement, volume, and cell-cell interactions) (Fig. 5A). The data generated can be directly imported into Flowjo for further analysis. Thus, Chrysalis and XTChrysalis2phtn automate histo-cytometry analysis of cell-cell interactions and protein expression in two-photon movies thereby reducing the required hands-on analysis time.

**Figure 5.**
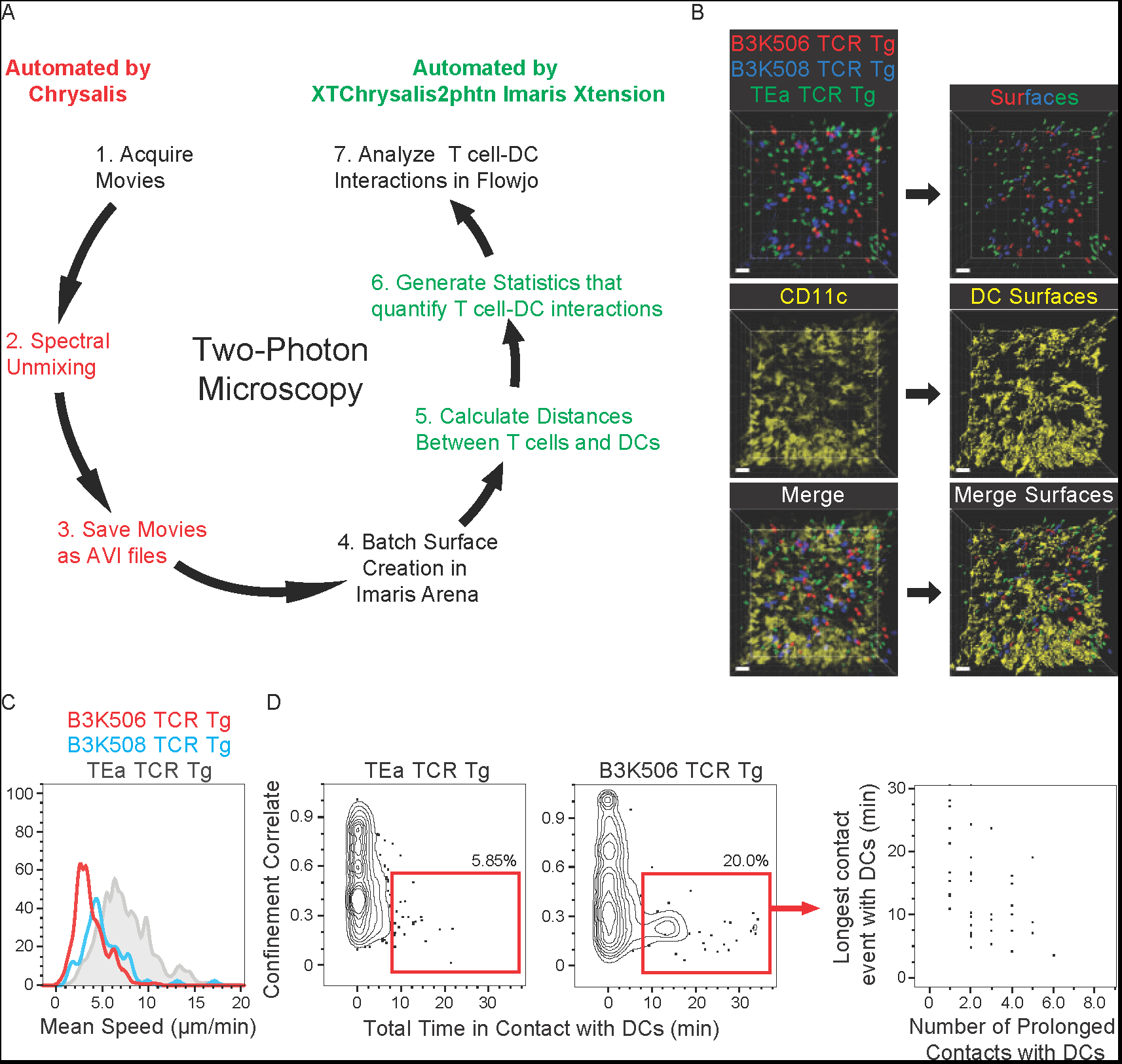
Chrysalis and XTChrysalis2phtn analysis of a two-photon microscopy movie. (A) Diagram of the histo-cytometry workflow on two-photon movies when automated by Chrysalis and XTChrysalis2phtn. (B) Surface-mediated identification of B3K506, B3K508, and TEa TCR Tg cells as well as DCs in two-photon movies. (C) Quantifying cellular velocity in a two-photon movie with Flowjo for B3K506 (red), B3K508 (blue), and TEa (grey) TCR Tg T cells. (D) Flowjo analysis of B3K506 and TEa TCR Tg cells in a two-photon movie, with quantification of track straightness, total contact time with DCs, longest contact with a DC, and number of prolonged contacts with DCs. Scale bar, 20 µm. Data representing two to three independent experiments are shown.

To demonstrate this improved workflow, T cell-DC interactions were quantified in two-photon microscopy movies depicting spleens from B6 recipients of B3K506, B3K508, and TEa TCR Tg cells infected 16 h earlier with *Lm*-P5R bacteria. The two-photon movies had 4 colors, which identified DCs and the three different TCR Tg populations (Fig. 5B) (19). Chrysalis spectrally unmixed and rescaled the movies, as well as generated AVI files to determine tissue health. For further analysis, the processed movies were opened in Imaris and surfaces were generated for the DCs and TCR Tg populations (Fig. 5B). XTChrysalis2phtn then generated cell statistics for analysis in Flowjo, which provided a way to compare B3K506 and B3K508 T cells recognizing P5R:I-A^b^ on DCs. TEa TCR Tg cells served as control cells because they do not respond to the infection (16, 17). The B3K506 and B3K508 cells had lower mean velocity than the TEa cells, suggesting that B3K506 and B3K508 cells interacted with DCs after infection while TEa cells did not (Fig. 5C). In line with this hypothesis, B3K506 cells had a lower confinement correlate value and greater contact time with DCs than TEa cells (Fig. 5D). Histo-cytometry analysis of these T cell-DC interactions allowed for a more granular view of these interactions by quantifying the duration of the longest contact event as well as the number of prolonged contact events for each T cell (Fig. 5D). As expected, T cells with the longest contact events with DCs made fewer total contacts with DCs (Fig. 5D). This example demonstrates a powerful and streamlined workflow for analyzing two-photon movies.

## Discussion

The Chrysalis software and Imaris Xtensions described in this manuscript can be applied to a broad range of biological questions, while reducing analysis time and empowering quantitative image analysis. We demonstrated the power of this workflow by quantifying T cell localization within splenic tissue in epi-fluorescence images, T cell-DC interactions in confocal microscopy images, and T cell motility and T cell-DC interactions in two-photon microscopy images. These same approaches can answer other immunological questions that require the quantification of cell localization, cell-cell interactions, or the ability to subset cells in images.

To extend the capabilities of this workflow beyond the applications described in this manuscript, we also generated separate Imaris Xtensions for each of the major steps performed by XTChrysalis, like batched statistics export. With these additional Xtensions, users can daisy chain Xtensions to batch image analysis in a manner that specifically addresses their research question. To further facilitate the use of this quantitative imaging approach in immunological research, we provide a step-by-step protocol that incorporates the automation steps detailed in this manuscript to streamline acquisition and analysis of confocal, epi-fluorescence, and two-photon microscopy images (Supplemental Protoc. 1).

While our protocol utilizes the commercial image analysis software Imaris, this protocol and the Chrysalis image processing software we developed can also be paired with free, publically available software such as CellProfiler and ilastik (51–54). While these programs do not have all of the features of Imaris, these programs are able to perform cell segmentation to identify cells within images, an essential step in the histo-cytometry workflow that is performed by Imaris in our protocol. Additionally, our protocol utilizes the commercial software Flowjo for comparing and quantifying image-derived statistics for each identified cell population. However, free, publicly available software can be used within our existing workflow in place of Flowjo for quantifying images, such as XiT and FACSanadu (26, 55).

To further reduce analysis time, we developed a signal absorption technique that expedites the quantification of cellular localization. The premise of this method is that a cell near other cells will absorb the nearby cell’s fluorescence. For example, a T cell residing in a B cell follicle will absorb B220 signal from nearby B cells. Signal absorption can then be used as a readout of cell location. This strategy is favorable over directly quantifying cell distance to a tissue structure because signal absorption only requires creating surfaces for cells and measuring their fluorescent signal. Conversely, the distance quantification approach involves creating surfaces for cells and tissue structures before quantifying the cells distance to the tissue structure. While the distance quantification approach provides a more definitive determination of localization, the extra steps of this approach require greater hands-on analysis time and computational power. This problem is especially exacerbated when the distance quantification approach is applied to the analysis of large tissues, like the spleen, or many biological samples, like during a time course. Therefore, the signal absorption strategy is a simpler and more time-efficient approach for quantifying cellular localization in these cases.

In summary, Chrysalis and the suite of Imaris Xtensions provide a high-throughput image processing workflow for confocal, epi-fluorescence, and two-photon microscopy images. This approach identifies subtle differences in cell phenotype and cell-cell interactions, while also offering significant reduction in hands-on analysis time. This time-savings reduces the barrier of entry for conducting quantitative, multispectral image analysis. Accessibility to this image analysis pipeline is further enhanced by the accompanying step-by-step protocol describing how to prepare samples, acquire images, and analyze images using the novel Chrysalis software and Imaris Xtensions for confocal, epi-fluorescence, and two-photon microscopy images (Supplemental Protoc. 1). An increase in the widespread adoption of these powerful, quantitative image analysis approaches will allow for novel and counterintuitive discoveries about the function and maintenance of the immune system.

## Acknowledgments

We thank J. Walter and C. Ellwood for technical assistance, and J. Kotov for reviewing the manuscript. We thank P. Beemiller for creating Sortomato, and M.Y. Gerner for helpful suggestions on histo-cytometry. D.I.K., T.P., J.S.M., and M.J.K. declare no competing interest. M.J.G. is employed by Bitplane, which produces the Imaris image analysis software that is utilized extensively in the image analysis pipeline described in this manuscript.

## Author Contributions

D.I.K. designed the study, wrote software, performed experiments, analyzed data, and wrote the manuscript, T.P. and M.J.G. wrote software, J.S.M. performed experiments, and M.K.J. critically reviewed the manuscript.

## Footnotes

This work was supported by grants to DIK (T32 AI83196 and T32 AI007313) and MKJ (R01 AI039614).

Non-standard Abbreviations:

dendritic cell (DC)

C57BL/6 (B6)

*Listeria monocytogenes* (*Lm*)

*Listeria monocytogene* expressing P5R (*Lm*-P5R)

MHC class II (MHCII)

transgenic (Tg)

phosphorylated form of S6 kinase (pS6)

Type 1 helper (Th1)

T follicular helper (Tfh)

P5R peptide bound to I-A^b^ (P5R:I-A^b^)

time-lapse data (movies)

